## Supplemental Tables for "Human influenza A virus H1N1 in marine mammals in California, 2019"

**Table 1: Metadata for IAV RNA positive marine mammals on the California Coast, 2016-2019.** The second sample on which ELISA or HI was performed upon was collected between 2 to 3 weeks after the admit sample. CSL is California sea lion, NES is northern elephant seal, NFS is northern fur seal, PHS is Pacific harbor seal. ND indicates no sample was available for testing, F is female, M is male, N/A indicates not applicable, Neg is negative, Pos is positive, HI is hemagglutination inhibition, ELISA is enzyme linked immunosorbent assay, RRT-PCR is real time reverse transcription polymerase chain reaction, Ct is cycle threshold. Asterisks indicate that HI was performed on the first sample since for some animals a second sample was not available. The number after each H designation indicates the hemagglutinin subtype used in the HI assay. A non-HI reactive sample is annotated at <8, where a 1:8 dilution was the lowest tested. Each RRT-PCR Ct and ELISA represent testing of a single replicate for each sample. For HI, each sample was tested in triplicate and the mean HI titer is shown.

| **Animal ID** | **IAV RRT-PCR Ct** | **ELISA** | | **Mean HI titer for second sample** | | | **Age Class** | **Sex** | **Species** | **Strand County** | **Strand City** | **Admit Date** | **Disposition** | | **Diagnosis** | **Cause of Death** |
| --- | --- | --- | --- | --- | --- | --- | --- | --- | --- | --- | --- | --- | --- | --- | --- | --- |
|  |  | **admit** | **second** | **H1** | **H3** | **H5** |  |  |  |  |  |  | **Date** | **Status** |  |  |
| CSL-13160 | 37.6 | Neg | ND | ND | ND | ND | Subadult | F | CSL | San Mateo | Pescadero | 06/05/2016 | 06/18/2016 | Died in Treatment | malnutrition | malnutrition |
| CSL-13161 | 42.0 | Neg | ND | ND | ND | ND | Adult | F | CSL | Santa Cruz | Watsonville | 06/06/2016 | 06/10/2016 | Euthanasia | N/A | euthanasia, domoic acid toxicity |
| CSL-13198 | 37.7 | Neg | ND | ND | ND | ND | Yearling | F | CSL | Monterey | Moss Landing | 07/04/2016 | 07/05/2016 | Euthanasia | N/A | gunshot |
| CSL-13203 | 36.8 | NA | ND | ND | ND | ND | Adult | F | CSL | San Luis Obispo | Morro Bay | 07/12/2016 | 07/31/2016 | Euthanasia | N/A | domoic acid toxicity |
| CSL-13207 | 37.1 | NA | ND | ND | ND | ND | Adult | F | CSL | Monterey | Pacific Grove | 07/14/2016 | 07/16/2016 | Died in Treatment | N/A | renal failure, neoplasia |
| ES-4068 | 36.5 | Neg | ND | ND | ND | ND | Pup | M | NES | Santa Cruz | Davenport | 01/24/2017 | 06/07/2017 | Released | malnutrition, maternal separation | N/A |
| ES-4121 | 35.8 | Neg | ND | ND | ND | ND | Pup | M | NES | Monterey | Pacific Grove | 03/20/2017 | 04/27/2017 | Released | malnutrition | N/A |
| NFS-435 | 35.7 | Neg | ND | ND | ND | ND | Pup | F | NFS | Monterey | Monterey | 12/21/2017 | 02/02/2018 | Released | malnutrition | N/A |
| ES-4254 | 35.4 | Neg | ND | ND | ND | ND | Pup | F | NES | Santa Cruz | Davenport | 01/22/2018 | 06/22/2018 | Released | maternal, separation, malnutrition, | N/A |
| HS-2754 | 35.3 | Neg | ND | ND | ND | ND | Pup | M | PHS | Marin | Bolinas | 02/14/2018 | 02/14/2018 | Died in Treatment | N/A | prematurity, maternal separation |
| ES-4311 | 36.5 | Pos | ND | 32* | 8* | <8* | Pup | M | NES | Monterey | Monterey | 03/26/2018 | 06/22/2018 | Released | malnutrition | N/A |
| CSL-14130 | 37.6 | Neg | ND | ND | ND | ND | Pup | M | CSL | San Mateo | Princeton-by-the-Sea | 12/17/2018 | 01/22/2019 | Released | malnutrition, maternal separation | N/A |
| ES-4424 | 35.8 | Pos | ND | ND | ND | ND | Pup | F | NES | San Luis Obispo | San Simeon | 3/12/2019 | 4/24/2019 | Released | malnutrition | N/A |
| ES-4506 | 20.9 | Neg | ND | ND | ND | ND | Pup | M | NES | San Luis Obispo | Avila Beach | 4/9/2019 | 7/17/2019 | Released | malnutrition, oil | N/A |
| ES-4507 | 28.6 | Pos | ND | 8* | <8* | <8* | Pup | M | NES | Santa Cruz | Live Oak | 4/9/2019 | 4/26/2019 | Euthanized | malnutrition, otostrongyliasis | otostrongyliasis |
| ES-4509 | 32.4 | Neg | Pos | 64 | 11 | <8 | Pup | F | NES | San Mateo | Pacifica | 4/10/2019 | 5/21/2019 | Released | malnutrition | N/A |
| ES-4523 | 34.7 | Pos | Pos | 128 | <8 | <8 | Pup | F | NES | Monterey | Pacific Grove | 4/14/2019 | 7/17/2019 | Released | malnutrition | N/A |
| ES-4527 | 32.6 | Neg | Pos | 256 | <8 | <8 | Pup | F | NES | Sonoma | Fort Ross | 4/15/2019 | 6/12/2019 | Released | malnutrition, otostrongyliasis, abscess | N/A |
| ES-4530 | 32.1 | Neg | Neg | ND | ND | ND | Pup | M | NES | San Mateo | Montara | 4/16/2019 | 6/1/2019 | Released | malnutrition, otostrongyliasis | N/A |
| ES-4538 | 28.2 | Pos | ND | 128* | <8 | <8 | Pup | F | NES | San Luis Obispo | San Simeon | 4/20/2019 | 6/12/2019 | Released | malnutrition, trauma, unknown | N/A |
| ES-4539 | 24.2 | Neg | Neg | >512 | <8 | <8 | Pup | F | NES | San Luis Obispo | Cayucos | 4/20/2019 | 5/2/2019 | Euthanized | malnutrition, trauma, unknown | congenital defect |
| HS-2859 | 32.0 | Neg | ND | ND | ND | ND | Pup | F | PHS | San Mateo | Pacifica | 4/27/2019 | 6/18/2019 | Released | maternal separation, malnutrition | N/A |

**Table 2: IAV antibody hemagglutination inhibition (HI) assays detections in ELISA-IAV reactive Northern elephant seals on California coasts in 2018-2019.** Animals shaded in grey also contained IAV RNA detectable by RRT-PCR; the same HI data for those animals is also reproduced in Table 2 for comparison with RNA values. Ferret serum from an animal that was experimentally inoculated with IAV H1N1 was used as a positive control. Positive control sera for H3N8 and H5N2 were not available. The negative control consisted of serum diluent.

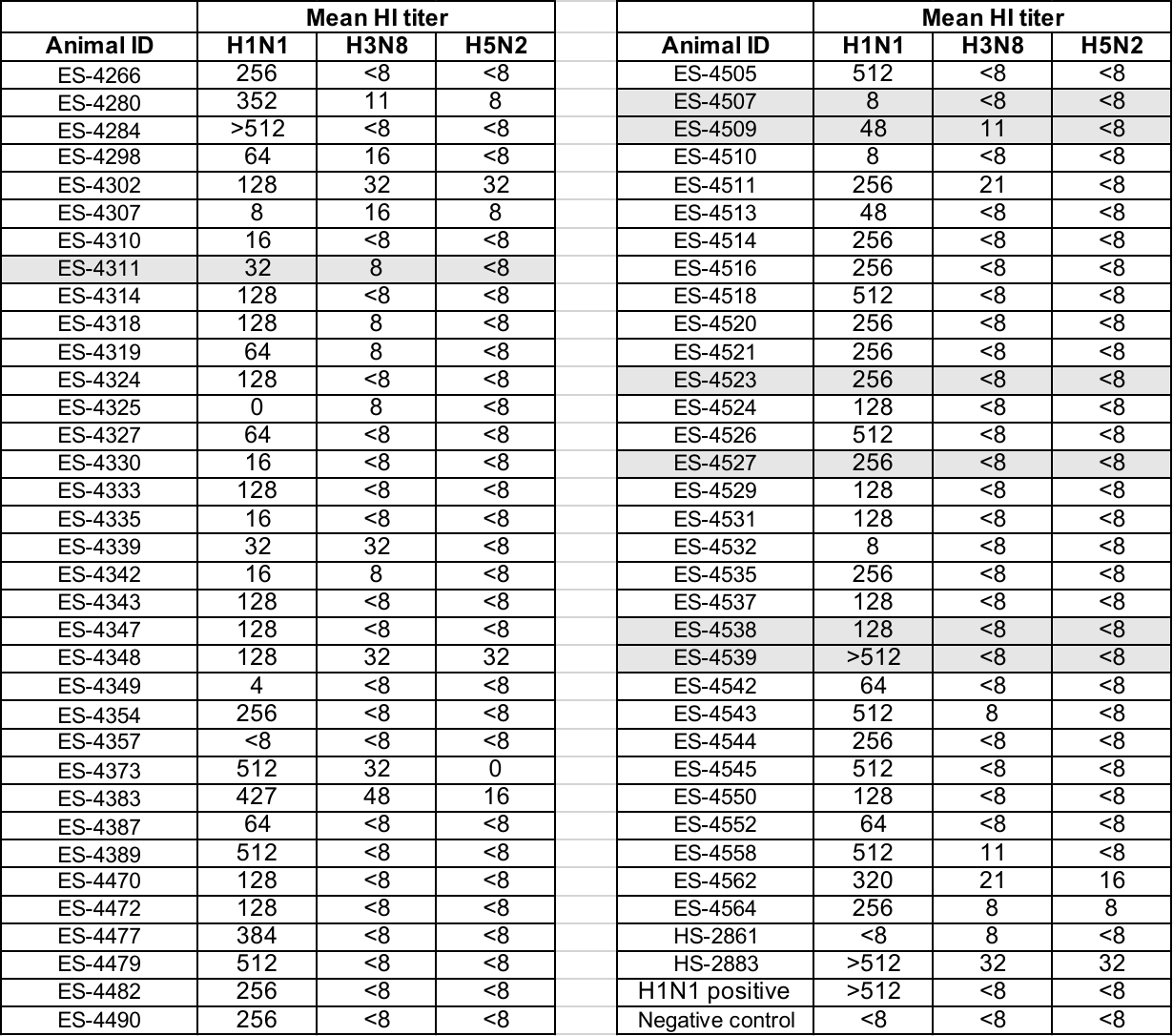
